# The NBSGW RIP-DTR Mouse: An Integrated Platform for Diabetes Induction, Human Immune Reconstitution and Transplantation Studies

**DOI:** 10.64898/2026.07.10.737744

**Authors:** Connie S. Chamberlain, Deep Kapadia, Ellen Abad Santos, Liupei Huang, Alexis M. Holm, Caterra Leavens, Ayesha Palwasha-Khan, Katherine M. Gorski, Colin C. Steck, Anna Mikat, Daniel M. Tremmel, Peter Chlebeck, Kayla Talerico, Esau Estrada, Brian E. McIntosh, Matthew E. Brown, Sara Dutton Sackett, Jon S. Odorico

## Abstract

Using toxin receptor-mediated cell ablation, a diabetes mouse model was generated that supports engraftment of human hematopoietic stem/progenitor cells (HSPCs) without the need for irradiation. The NBSGW immunodeficient strain was crossed with the NSG RIP-DTR which carries the diphtheria toxin receptor (hDTR) under the control of the rat insulin promoter to generate the NBSGW RIP-DTR mouse. This model enables controlled β-cell ablation, robust human immune system reconstitution without myeloablative conditioning, and evaluation of human immune-mediated graft rejection within a single platform. NBSGW RIP-DTR mice exhibited reproducible and titratable diabetes induction, supported durable human islet engraftment and glycemic correction, and retained efficient human hematopoietic reconstitution comparable to the parental NBSGW strain. In humanized mice, diphtheria toxin-mediated diabetes induction was well tolerated and enabled assessment of human immune responses to allogeneic islet grafts. Collectively, these findings establish the NBSGW RIP-DTR mouse as an integrated and clinically relevant platform for studying β-cell replacement therapies and human immune-mediated graft rejection.

**Article Highlights:**

- Current preclinical models do not simultaneously support controlled diabetes induction, human islet transplantation, and durable human immune reconstitution without irradiation.
- This study asked whether the NBSGW RIP-DTR mouse could integrate diphtheria toxin-mediated β-cell ablation with irradiation-free humanization in a single platform.
- NBSGW RIP-DTR mice demonstrated reproducible diabetes induction, supported functional human islet engraftment, and retained robust human hematopoietic reconstitution comparable to parental strains.
- These findings establish the NBSGW RIP-DTR model as a clinically relevant platform for studying β-cell replacement and human immune response to islet allografts, xenografts and stem cell-derived islets.

---

Preclinical models are central to studying β-cell replacement (e.g., via cadaveric islet or pluripotent stem cell [PSC]-derived islet transplant) and immune-mediated graft responses, yet existing systems remain fragmented. In practice, diabetes induction, human immune system reconstitution, and transplantation studies are often performed in separate models, limiting the ability to interrogate these processes within a unified experimental framework. This lack of integration complicates mechanistic interpretation and reduces translational relevance.

Humanized mouse models provide a means to study human immune responses *in vivo* but typically require myeloablative conditioning, which introduces toxicity, disrupts tissue homeostasis, and limits experimental duration (for reviews, see ^1, 2^). In addition, variability in human immune reconstitution can further limit reproducibility. Independently, commonly used approaches for diabetes induction, such as streptozotocin (STZ) ^3–5^, are associated with off-target cytotoxicity, variable penetrance, and incomplete β-cell ablation, complicating downstream analysis (for review see ^6^).

Genetic models based on the NOD.Cg-*Prkdc^scid^Il2rg^tm1Wjl^/*SzJ (NSG) background^7–9^ in which expression of the human diphtheria toxin receptor (DTR) is driven by the rat insulin promoter (RIP), enables selective and titratable β-cell ablation after diphtheria toxin (DT) administration.^10–12^ These RIP-DTR systems have been widely used to achieve targeted β-cell ablation and support studies of islet replacement and transplant immunology.^13^ However, the models typically rely on irradiation-based humanization strategies^14^, which limits their utility for studies requiring stable, long-term immune reconstitution.

The NBSGW strain, which carries the Kit*^W^*^41^ mutation, supports human hematopoietic engraftment without need for irradiation or chemical myeloablation to achieve human chimerism.^15, 16^ This feature provides an opportunity to integrate controlled β-cell ablation with irradiation-independent humanization within a single genetic background.

To address the paucity of preclinical models capable of simultaneously modeling diabetes, human immune reconstitution, and islet transplantation, we generated an NBSGW RIP-DTR mouse model by introducing the RIP-DTR transgene into the NBSGW background. This platform enables controlled diabetes induction, supports human immune reconstitution, and facilitates evaluation of transplanted islets within a single experimental system.

## RESEARCH DESIGN AND METHODS

### Mouse Models and Genotyping

All animal procedures were approved by the Institutional Animal Care and Use Committee (IACUC) at the University of Wisconsin-Madison and conducted in accordance with institutional guidelines. Mice were housed under specific pathogen-free conditions with ad libitum access to food and water on a 12-h light/dark cycle. Both male and female mice were included, and sex was recorded for all experiments. Sex-stratified analyses were performed when biologically relevant or sex-associated differences were observed. Sample sizes for each experiment are provided in the corresponding figure legends and tables.

NBSGW RIP-DTR mice were generated by crossing NSG RIP-DTR mice with NBSGW mice for 10 generations to establish a congenic strain (**Fig. 1A)**. Offspring carrying both the RIP-DTR transgene and the *Kit^W^*^41^ allele were identified and subsequently backcrossed to the NBSGW background to establish a stable strain. All integral genes were genotyped via TaqMan qPCR assay with specific probes designed for each specific gene of interest (Transnetyx, Cordova, TN). Experimental mice were required to carry the RIP-DTR transgene and the genetic background necessary to support immunodeficiency and human cell engraftment prior to inclusion in downstream studies.

**Figure 1.**
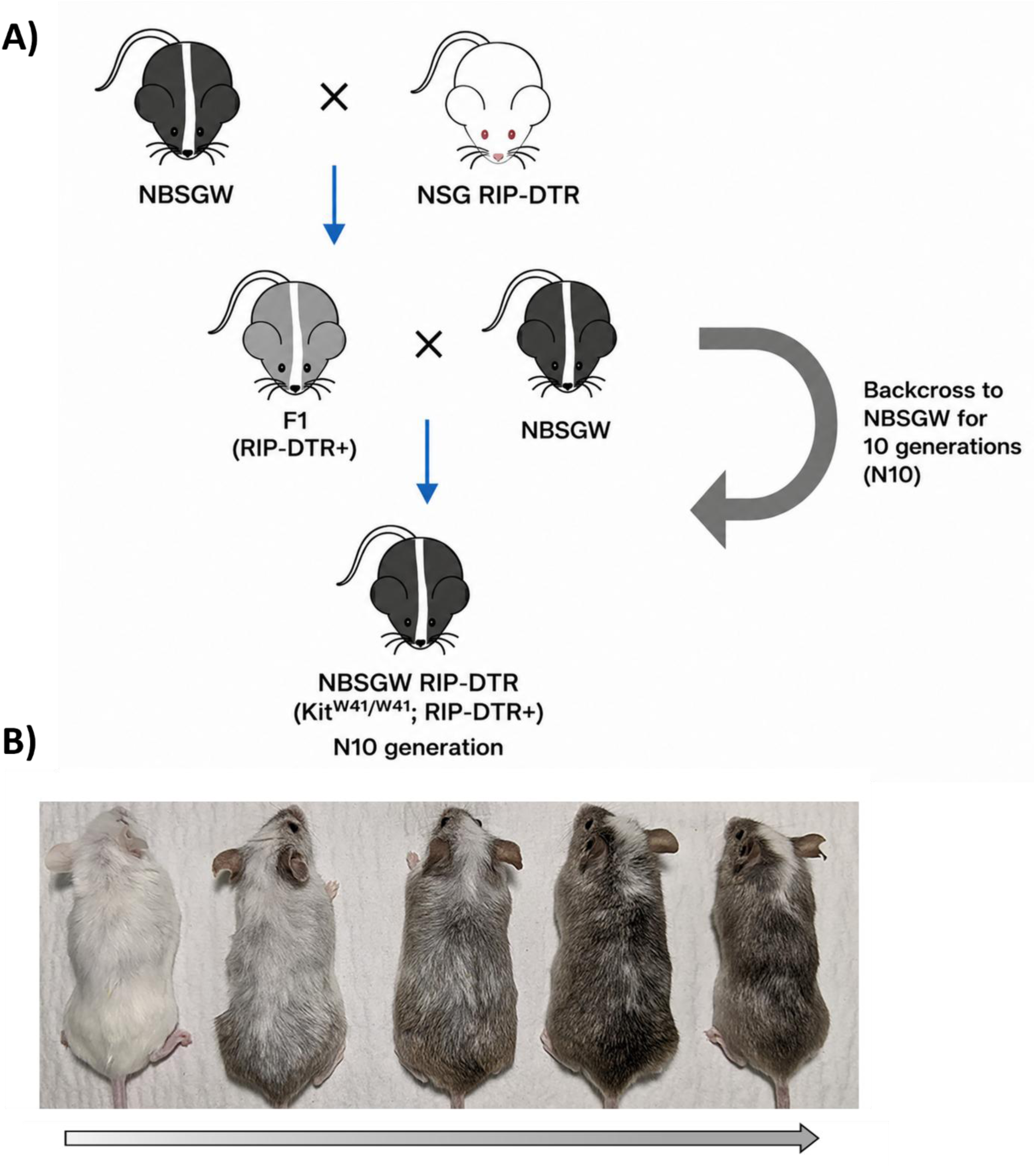
Generation of the NBSGW RIP-DTR mouse strain and visual confirmation of strain-associated coat color. **A)** Schematic illustrating the breeding strategy used to generate the NBSGW RIP-DTR strain. NBSGW mice were crossed with NSG RIP-DTR mice to generate F1 progeny carrying the RIP-DTR (*Ins2-DTR)* transgene. Transgene-positive offspring were subsequently backcrossed to NBSGW mice for 10 generations to establish the NBSGW RIP-DTR strain with a homozygous *Kit^W^*^41^*^/W^*^41^ NBSGW background while retaining the RIP-DTR transgene (N10 generation). **B)** Representative images of mice from successive breeding generations, showing the transition to the albino coat color characteristic of the NBSGW strain background. Images are representative of more than three independent litters.

Humanized mice were generated using the NeoThy approach, which combines transplantation of human neonatal thymic tissue fragments implanted under the renal capsule with autologous or allogeneic human cord blood derived CD34^+^ hematopoietic stem/progenitor cells (HSPCs), following anti-CD2 antibody treatment^17–19^. Non-fetal human tissues were obtained from approved sources in compliance with UW-Madison Health Sciences Institutional Review Board and federal guidelines. Mice were humanized without irradiation as previously reported.^15, 18,20^ Engraftment was assessed at 8-, 12-, and 15-16-weeks using flow cytometry. All animal work was conducted under approval of the UW-Madison Institutional Animal Care and Use Committee. Additional information is provided in the Supplemental Materials.

### Diabetes Induction and Glucose Monitoring

To identify DT regimens that reliably induced hyperglycemia while minimizing toxicity, an initial dose-ranging study was performed in NSG RIP-DTR mice. Based on these experiments, two different regimens of DT administration were selected for subsequent evaluation in NBSGW RIP-DTR mice: 5 ng DT intraperitoneal (i.p.) daily for three consecutive days (5 ng x 3) or given as a single bolus of 15 ng DT i.p. (15 ng x 1). Diabetes durability was monitored for up to 6 weeks, and insulin supplementation was provided as needed to maintain animal welfare. As a comparator, streptozotocin (STZ) was freshly prepared in cold citrate buffer (pH 4.5) and administered as a single i.p. dose of 150 mg/kg. Blood glucose (BG) and survival were monitored following administration.

Non-fasting BG was measured via tail-vein blood using a Contour Next glucometer. For DT dose-optimization studies, mice were considered hyperglycemic when BG exceeded ≥300 mg/dL. For transplantation studies, mice were considered diabetic and eligible for transplantation after BG remained ≥300 mg/dL for three consecutive measurements prior to transplantation. Additional information is provided in the Supplemental Materials.

### Stem Cell-Derived Islet (SCI) Differentiation

HUES8 human embryonic stem cells (hESCs) were differentiated into SCI using previously published six stage suspension differentiation protocol described by Hogrebe et al.^21^ Cells were sequentially differentiated through definitive endoderm, primitive gut tube, pancreatic progenitors 1/2, endocrine progenitor and β-cell maturation stages using stage-specific media supplemented with defined growth factors and small molecules. SCI were collected upon completion of Stage 6 and transplanted under the renal capsule of recipient mice. Additional information is provided in the Supplemental Materials.

### Human Islet Transplantation and Metabolic Assessment

Human pancreatic islets were provided by the NIDDK-funded Integrated Islet Distribution Program (IIDP) Donor characteristics, including age, sex, BMI, HbA1C, islet purity, viability, pre-transplant culture duration, and donor/source identifiers, are provided in **Supplementary Table 1.** Islets were cultured in PIMR at 37°C/5%CO_2_ before transplantation and were used within 7 days after isolation.

At the time of transplantation, mice underwent renal subcapsular (KSC) transplantation as previously described^22^. Briefly, mice received either 2,000 human pancreatic islet equivalents (IEQ) or 10,000 IEQ SCIs under isoflurane anesthesia. Mice were monitored for up to 12 weeks after transplantation, with longitudinal assessment of BG and graft function. To maintain animal welfare during periods of hyperglycemia, exogenous insulin was administered via SQ insulin glargine injection or LinBit sustained-release insulin implants.

Peripheral blood was collected before transplantation and every 2 weeks for measurement of circulating human C-peptide as an indicator of graft function. Fed BG was monitored twice weekly throughout the study.

At 12 weeks post-transplantation, mice underwent intraperitoneal glucose tolerance testing (IPGTT) following an overnight fast. Glucose (2g/kg BW) was administered i.p., and BG concentrations were measured serially to assess dynamic glucose regulation. Glucose excursion/disposal was quantified by calculating the area under the curve (AUC). Additional methodological details are provided in the Supplemental Materials.

### Histology, Immunofluorescence, and Imaging

Kidney grafts were harvested at study endpoint and processed for histological and immunofluorescence analysis.

Hematoxylin and eosin (H&E) staining was performed to assess graft morphology. Staining for insulin and glucagon was performed to assess endocrine cell composition and graft integrity. Staining for CD3, CD45, and CD68 was performed to assess immune cell infiltration and graft rejection. Antibodies used are listed in **Supplementary Table 2**.

Fluorescence and H&E images were acquired using standard microscopy platforms. Images were collected using identical acquisition setting within each experiment and saved as uncompressed TIFF files. Brightness and contrast adjustments, if applied, were performed uniformly across comparable images. Additional information is provided in the Supplemental Materials.

### Flow Cytometric Analysis of Human Engraftment

Flow cytometry was used to quantify human immune cell engraftment and lymphocyte subsets and was performed as previously reported.^15, 18, 20^ Human CD45^+^ cells were identified and used to calculate overall engraftment. Subsets of T and B cells were analyzed within the human leukocyte population. Antibodies used are listed in **Supplementary Table 2.**

### Statistical Analysis

Data were analyzed using linear mixed-effects models to account for repeated measurements within animals. Variance components were estimated using restricted maximum likelihood (REML). Post hoc pairwise comparisons were adjusted using Tukey’s multiple-comparison test. Time-to-event outcomes were analyzed by Kaplan-Meier analysis with log-rank testing. Statistical significance was defined as p < 0.05. All statistical analyses were performed in GraphPad Prism, v10 (GraphPad Software, Boston, MA).

## RESULTS

### Generation and validation of NBSGW RIP-DTR mice

PCR-based genotyping confirmed that all animals used for subsequent experiments carried the required allelic combination **(Table 1).** Mice were homozygous for the Kit*^W^*^41^*^/W^*^41^ mutation and retained the *Prkdc^scid^* and *Il2rg-null* alleles required for immunodeficiency and human immune cell engraftment (**Supplementary Fig. 1A**). All animals were also positive for the Ins2-DTR (human *HB-EGF*) transgene, enabling β-cell specific sensitivity to DT (**Supplementary Fig. 1B)**.

**Table 1.** Genotype verification of NBSGW RIP-DTR mice.

| Genetic Element | Gene /Locus | Expected Genotype | Purpose of Assay |
| --- | --- | --- | --- |
| Kit W41 mutation | *Kit* | Kit<sup>W41/W41</sup> | Confirms homozygosity of the Kit<sup>W41</sup> allele, a defining feature of the NBSGW background that enables efficient human hematopoietic engraftment without irradiation. |
| SCID mutation | *Prkdc* | *Prkdc*<sup>scid/scid</sup> | Confirms the *Prkdc*<sup>scid/scid</sup> responsible for severe combined immunodeficiency and loss of adaptive immune function. |
| *Il2rg* knockout | *Il2rg* | Null allele present | Confirms the *Il2rg* null allele required for absence of NK cells and enhanced human cell engraftment. |
| Ins2-DTR transgene | *Ins2-HB-EGF* | Transgene positive | Confirms the RIP-DTR transgene, which places the diphtheria toxin receptor (DTR) under the control of the rat insulin promoter (RIP), allowing selective ablation of β-cells after diphtheria toxin treatment. |
| Tyrosinase mutation | *Tyr* | Mutant | Confirms the albino *Tyr* mutation associated with the NBSGW background. |

Progressive coat color uniformity consistent with the NBSGW phenotype was observed across breeding generations **(Fig. 1B)**, supporting successful strain stabilization. Together, these findings confirm successful generation of the NBSGW RIP-DTR model.

### DT-mediated diabetes induction is preserved in the NBSGW RIP-DTR background

Dose-ranging studies in NSG RIP-DTR mice identified two regimens, 5 ng x 3 and 15 ng x 1, that produced rapid and reproducible diabetes with 100% hyperglycemia by day 7 (**Supplementary Table 3)**. In contrast, the 10 ng x 1 regimen was suboptimal, resulting in delayed onset and reduced diabetes penetrance. Increasing the DT dose beyond 30 ng did not improve penetrance or accelerate disease onset, indicating a plateau effect. However, higher DT doses produced more severe hyperglycemia, with 55.2% of mice reaching the upper detection limit of the glucometer (600 mg/dL), compared with 9.9% of mice receiving the selected lower-dose regimens. Sex-stratified analysis revealed modest differences in onset but similar overall penetrance.

The 5 ng x 3 and 15 ng x 1 regimens were therefore selected for evaluation in NBSGW RIP-DTR mice (**Fig 2A)**. Following DT administration, NBSGW RIP-DTR and NSG RIP-DTR mice developed comparable progressive hyperglycemia with no overall strain effect (p = 0.28), indicating that introduction of the NBSGW background to the NSG RIP-DTR model did not impair the magnitude of DT-induced hyperglycemia (**Fig. 2B**). Under the 5 ng x 3 dosing regimen, hyperglycemia developed in 100% of NBSGW RIP-DTR mice and 97.4% of NSG RIP-DTR mice, with median onset occurring on days 4 and 3, respectively (**Fig. 2C**). Longitudinal analysis demonstrated no overall strain-dependent differences (**Fig. 2D**). The single-dose 15 ng x 1 regimen also induced hyperglycemia efficiently in both strains, with 95.2% of NBSGW RIP-DTR mice and 84.6% of NSG RIP-DTR mice becoming hyperglycemic by day 13 (**Fig. 2C)**. BG increased significantly over time (p < 0.01), with no overall strain effect (**Fig 2E**). Representative pancreatic immunofluorescence confirmed marked loss of insulin-positive β-cells following DT administration in NBSGW RIP-DTR mice (**Fig. 2F)**. To determine whether DT-induced diabetes was durable, BG was monitored for up to six weeks following DT administration. BG remained elevated over time in both strains following induction, with no consistent strain-dependent differences **(Fig. 2G)**. Sex-stratified analysis demonstrated comparable responses to DT in NSG RIP-DTR and NBSGW RIP-DTR mice, with no meaningful strain-dependent differences in diabetes induction (**Supplementary Figs. 2A-B)**. Although males exhibited a modestly earlier increase in BG than females; diabetes penetrance and the subsequent glycemic course were similar between sexes.

**Figure 2.**
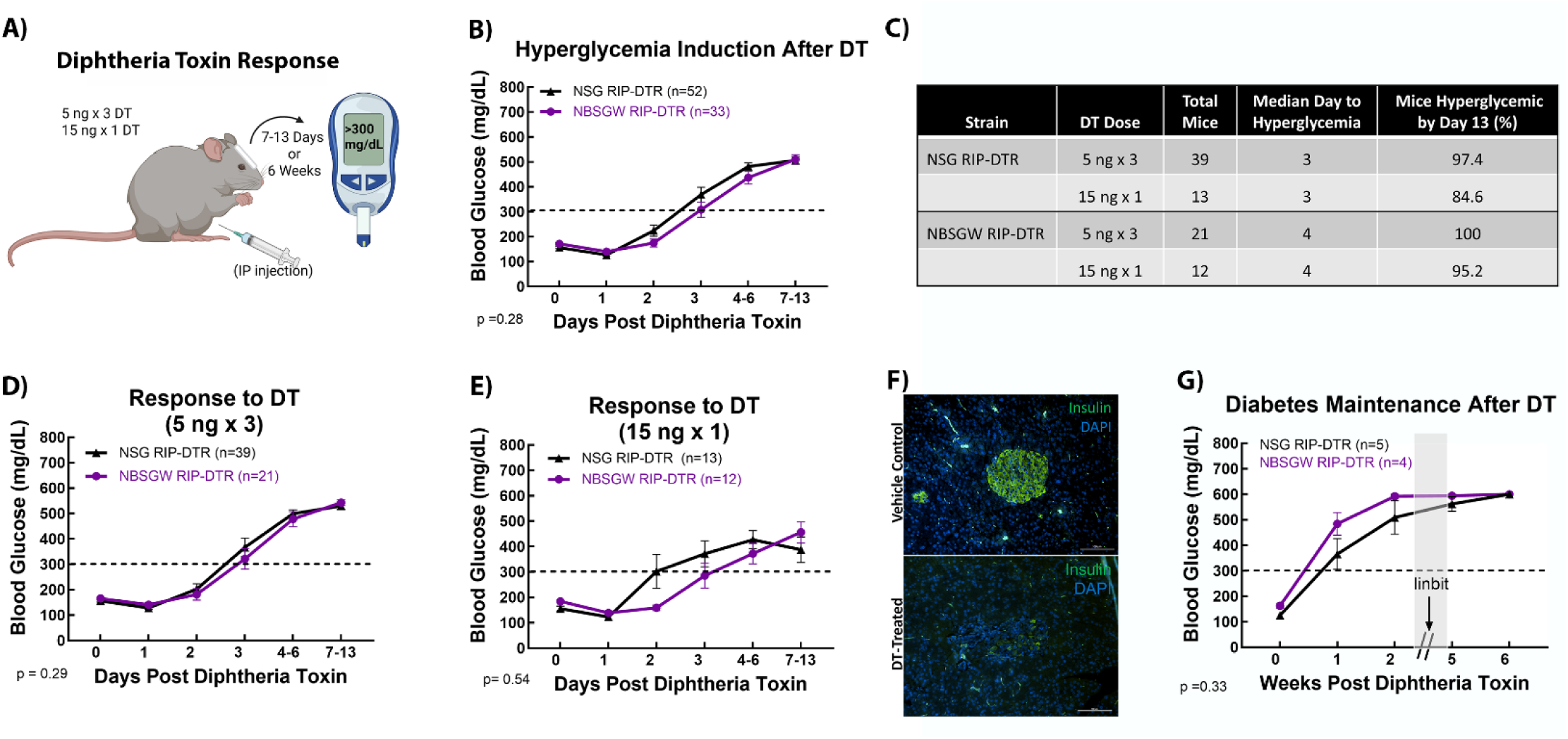
Diabetes induction of NBSGW RIP-DTR and NSG RIP-DTR mice following diphtheria toxin (DT) administration. **A)** Experimental schematic illustrating DT dosing regimens (5 ng administered on three consecutive days [5ng x 3] or a single 15 ng dose [15 ng x 1]) delivered via intraperitoneal injection, followed by blood glucose (BG) monitoring. Mice were monitored for 7-13 days for hyperglycemia induction studies or up to 6 weeks for long-term assessment. **B)** Longitudinal BG measurements following DT administration. Both strains exhibited rapid increases in BG, reaching hyperglycemic levels within 4-6 days. **C)** Summary of hyperglycemia induction by DT regimen, including the median day to hyperglycemia and the percentage of mice that developed hyperglycemia by day 13. **D-E)** BG trajectories following DT treatment regimens of either: **D)** 5 ng x 3 or **E)** 15 ng x 1. Both regimens effectively induced hyperglycemia in NBSGW RIP-DTR and NSG RIP-DTR mice. **F)** Representative pancreatic immunofluorescence images of NBSGW RIP-DTR mice. Insulin-positive β-cells (green) are present in vehicle-treated (Vehicle Control) pancreata. DT administration (DT-Treated) resulted in marked loss of insulin-positive β-cells. DAPI-stained nuclei are shown in blue. **G)** Sustained hyperglycemia following DT administration. Hyperglycemia persisted throughout the 6-week study following DT administration in both strains. A LinBit sustained-release insulin pellet was implanted after week 2 (arrow) to support animal welfare during prolonged hyperglycemia. The interval (grey shaded region) during which BG was transiently reduced by LinBit treatment is omitted for clarity. Data are presented as mean ± SEM. Statistical analysis was performed using mixed-effects models (REML) with fixed effects for strain, time, and DT dose, followed by Tukey’s post hoc testing (p< 0.05). Hyperglycemia was defined as BG ≥300 mg/dL (dashed line)

### NBSGW RIP-DTR mice support human islet-mediated reversal of diabetes

To determine whether introduction of the NBSGW background affects the capacity to support functional human islet transplantation, diabetes was induced in NBSGW RIP-DTR and NSG RIP-DTR mice before transplantation of 2,000 human IEQ beneath the renal capsule. (**Fig. 3A)**. Because biological sex influences susceptibility to experimental diabetes and glucose homeostasis in mice, this analysis focused on female recipients to enable direct comparison between strains.^23, 24^ Following transplantation, female recipients from both strains showed rapid improvement in fed BG, consistent with functional human islet engraftment (**Fig. 3B)**. Most animals achieved BG levels below 250 mg/dL within the first week after transplantation and maintained improved glycemic control throughout the 12-week observation period (**Fig. 3B**). Time to sustained normoglycemia was defined as the first occurrence of BG < 250 mg/dL. Kaplan-Meier analysis demonstrated comparable reversal kinetics between NBSGW RIP-DTR and NSG RIP-DTR female recipients (p = 0.56; **Fig. 3C**). Although NSG RIP-DTR females showed a slight trend toward earlier reversal, the overall frequency and durability of glycemic correction were similar between strains.

**Figure 3.**
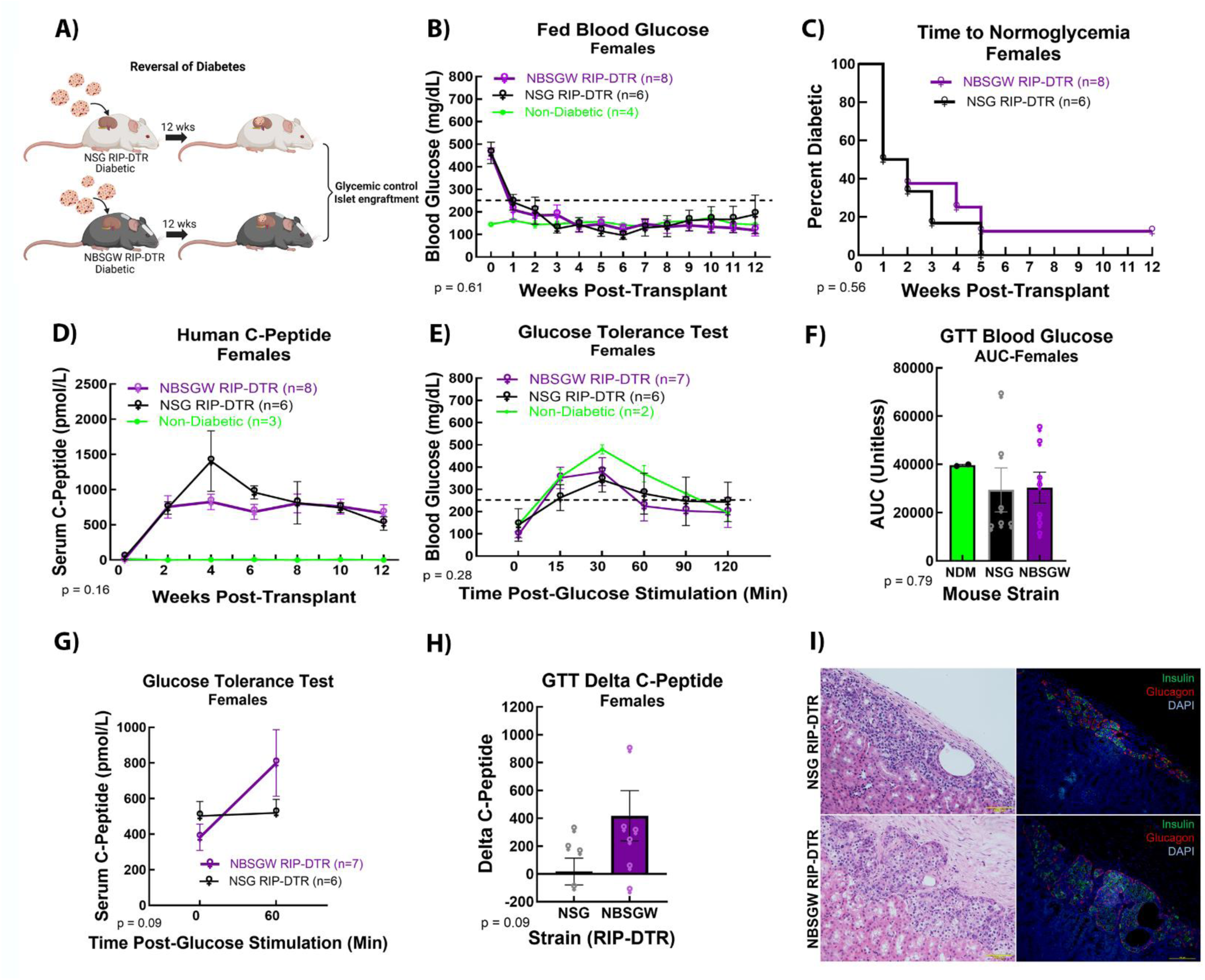
Female NBSGW RIP-DTR mice support reversal of diabetes following human islet transplantation. **A)** Experimental schematic of diabetes induction, human islet transplantation, and longitudinal metabolic assessment. **B)** Longitudinal fed BG levels following islet transplantation of 2000 human IEQ. **C)** Kaplan-Meier analysis of time to sustained normoglycemia defined as the first occurrence of BG <250 mg/dL. **D)** Fed serum human C-peptide levels following transplantation, demonstrating sustained graft function in both recipient strains. **E)** Intraperitoneal glucose tolerance testing (IPGTT) performed 12 weeks post-transplant to fasted mice. **F)** Area under the curve (AUC) for the IPGTT. **G)** Human C-peptide concentrations measured at baseline (0 min) and 60 min following glucose administration. **H)** Change in human C-peptide (Delta C-peptide; 60 min-baseline) during the IPGTT. **I)** Representative H&E (**left**) and immunofluorescence (**right**) images of human islet grafts beneath the renal capsule in the NSG RIP-DTR (**top**) and NBSGW RIP-DTR (**bottom**) mice. Insulin (**green**) and glucagon (**red**) staining with DAPI (**blue**), reveal preserved endocrine cell composition and comparable islet architecture between the two strains. Data are presented as mean ± SEM. Statistical analyses were performed using REML with Tukey’s post-hoc correction. For figures B,D,E,and F, the non-diabetic control group (green) is included as a reference and was not included in the analysis. The dotted line indicates the prespecified glycemic reversal threshold of BG <250 mg/dL.

Functional graft activity was confirmed by circulating human fed C-peptide (**Fig. 3D)**. C-peptide increased after transplantation in both groups and remained detectable over time, indicating that the NBSGW background does not impair human islet engraftment or sustained graft function compared with NSG RIP-DTR females.

At 12 weeks post-transplantation, intraperitoneal glucose tolerance testing (IPGTT) was performed in fasted mice to assess dynamic graft function. Following glucose challenge, both NBSGW RIP-DTR and NSG RIP-DTR mice exhibited a rapid rise in BG that peaked between 15-30 minutes, followed by a gradual decline toward baseline by 120 minutes (**Fig. 3E)**. Quantification of glucose excursion by area under the curve (AUC) revealed no significant difference between strains (**Fig. 3F).** Glucose stimulation also elicited increases in circulating human C-peptide in both groups (**Fig. 3G**). Although not statistically significant, NBSGW RIP-DTR females showed a modest trend toward greater C-peptide release at 60 minutes after glucose challenge. Change in C-peptide, was similar between strains (**Fig. 3H),** confirming preserved glucose-responsive insulin secretion.

Histological analysis of islet grafts at 12 weeks post-transplantation confirmed durable graft persistence and endocrine composition **(Fig. 3I)**. H&E staining demonstrated intact islets beneath the renal capsule. Immunofluorescence staining for insulin and glucagon confirmed maintenance of β- and α-cell populations within the graft, with comparable graft architecture between NSG RIP-DTR and NBSGW RIP-DTR mice.

Overall, these results demonstrate that female NBSGW RIP-DTR mice support robust human islet engraftment, durable glycemic correction, and physiologically responsive insulin secretion at levels comparable to that observed in NSG RIP-DTR mice.

### RIP-DTR expression does not impair irradiation-free human immune reconstitution

To determine whether introduction of the RIP-DTR transgene affects the ability of NBSGW mice to support irradiation-free human immune reconstitution, NeoThy-humanized NBSGW RIP-DTR and parental NBSGW mice were compared longitudinally following immune reconstitution (**Fig. 4A**). Peripheral blood was analyzed at 8-, 12-, and 15-16-weeks post-engraftment by flow cytometry to quantify human (h) CD45^+^ leukocytes, hCD19^+^ B cells, and hCD3^+^ T cells.

**Figure 4.**
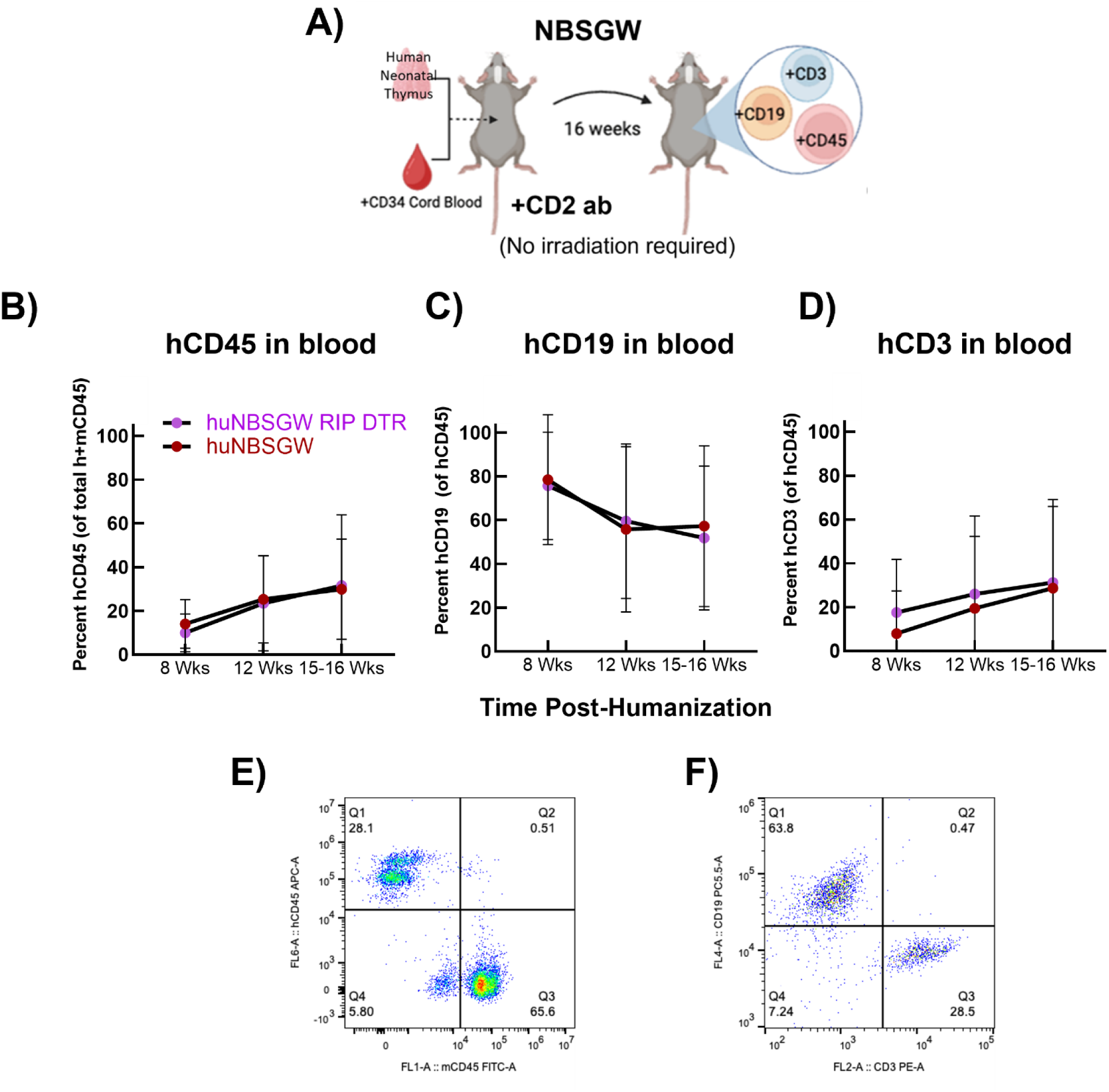
Human immune reconstitution in NBSGW RIP-DTR and NBSGW mice. **A)** Experimental schematic illustrating human immune reconstitution of NBSGW and NBSGW RIP-DTR mice by transplantation of human neonatal thymus tissue and CD34^+^ umbilical cord blood cells following anti-CD2 antibody treatment, without irradiation. **B-D)** Longitudinal frequencies of human immune cell population in peripheral blood at 8, 12, and 15-16 weeks following immune reconstitution, as determined via flow cytometry: **B)** human (h) CD45^+^ leukocytes, **C)** hCD19^+^ B cells, **D)** and hCD3^+^ T-cells. **E-F)** Representative flow cytometry plots illustrating gating strategy for **E)** hCD45^+^ leukocytes and **F)** hCD19^+^ and hCD3^+^ lymphocyte populations. Data are presented as mean ± SEM. Statistical comparisons were performed using REML with Tukey’s post-hoc correction. No significant differences were observed between humanized (hu) NBSGW RIP-DTR and huNBSGW mice for hCD45^+^ (p=0.98), hCD19^+^ (p=0.67) and hCD3^+^ (p=0.29).

Across all time points, NBSGW RIP-DTR mice exhibited hCD45^+^ leukocyte frequencies comparable to those observed in parental NBSGW controls, indicating that introduction of the RIP-DTR allele does not impair hematopoietic engraftment (**Fig. 4B**). Similarly, the frequencies of hCD19^+^ B cells (**Fig. 4C**) and hCD3^+^ T cells (**Fig. 4D**) within the hCD45^+^ leukocyte compartment were comparable between strains, indicating preserved lymphoid lineage development. Representative flow cytometry plots confirmed robust hCD45^+^ engraftment and identification of hCD19^+^ B cell and hCD3^+^ T cell populations in both strains (**Figs. 4E-F**). Together, these findings indicate that introduction of the RIP-DTR transgene does not alter the level or composition of human immune reconstitution in NBSGW mice.

### DT safely induces diabetes in humanized NBSGW RIP-DTR mice

To determine whether human immune reconstitution alters the efficiency of DT-mediated diabetes induction, female huNBSGW RIP-DTR mice were compared with non-humanized NSG RIP-DTR and NBSGW RIP-DTR mice following DT administration (**Fig. 5A).** Following DT administration, BG increased over time in all three groups (**Fig. 5B**). Hyperglycemia developed in 71.4% of huNBSGW RIP-DTR, 87.5% in NBSGW RIP-DTR and 80% in NSG RIP-DTR mice (**Fig. 5C**). The proportion of mice that developed hyperglycemia did not differ significantly among groups, indicating that DT induced diabetes with comparable efficiency in humanized and non-humanized female RIP-DTR mice.

**Figure 5.**
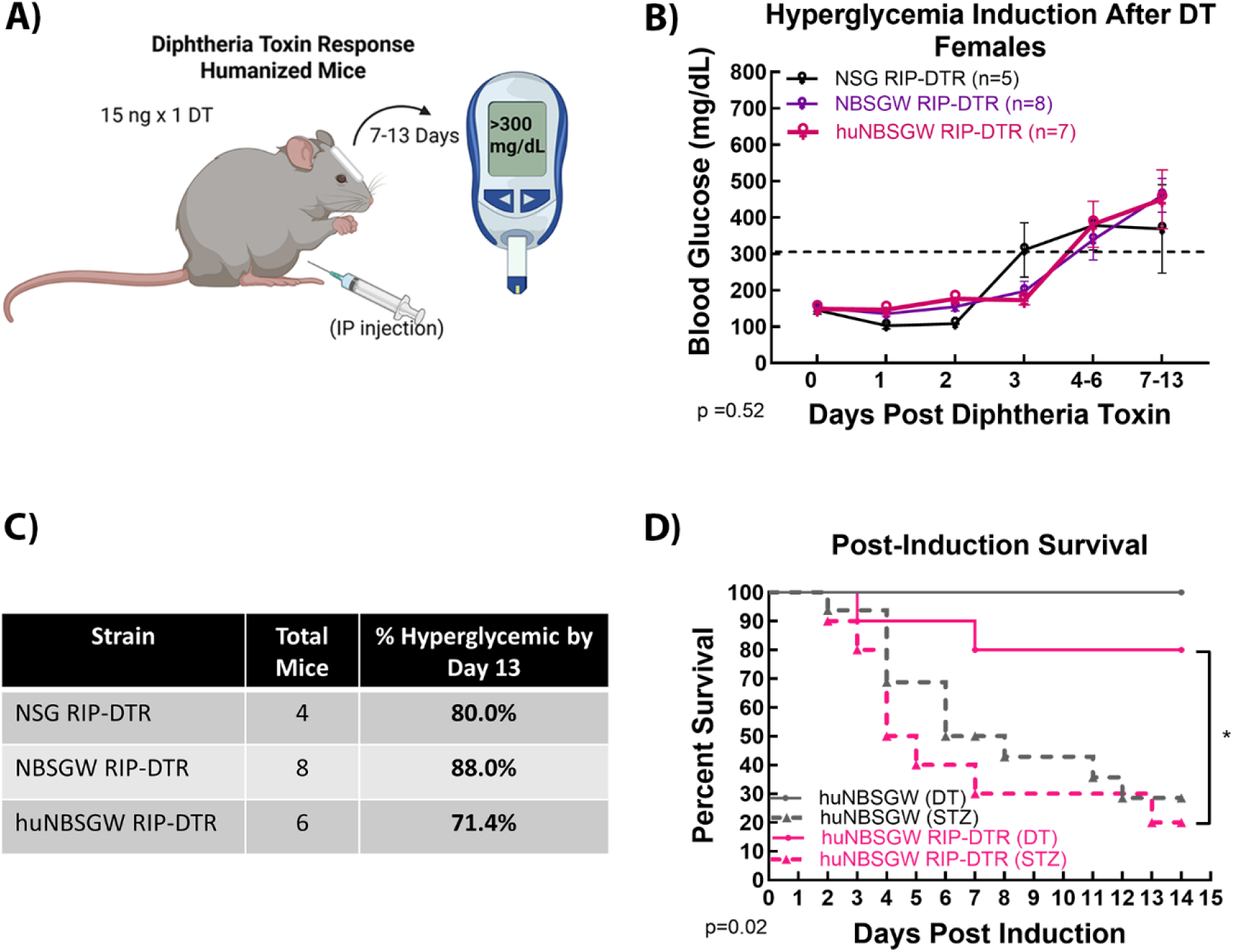
Diabetes induction of huNBSGW RIP-DTR mice. **A)** Experimental schematic illustrating diabetes induction by a single i.p. injection of 15 ng DT (15 ng x 1) and the subsequent monitoring period for hyperglycemia. **B)** Longitudinal fed BG measurement in female NSG RIP-DTR, NBSGW RIP-DTR, and humanized (hu) NBSGW RIP-DTR mice following DT administration. The dotted line indicates the hyperglycemia threshold (BG ≥300 mg/dL). **C)** Summary of diabetes incidence, presented as the percentage of mice that developed hyperglycemia by day 13 following DT (15 ng x 1) administration. **D)** Kaplan-Meier survival analysis of huNBSGW RIP-DTR and huNBSGW mice following diabetes induction with DT (n=10 huNBSGW RIP-DTR; n=2 huNBSGW) or streptozotocin (STZ; n=10 huNBSGW RIP-DTR; n=16 huNBSGW). Data are presented as mean ± SEM. Statistical comparisons were performed using REML with Tukey’s post-hoc correction **(B)**. Survival curves were analyzed using the Kaplan-Meier method with log-rank (Mantel-Cox) test **(D)**. Statistical significance was based on p<0.05. *p < 0.05.

Notably, DT administration was well-tolerated in huNBSGW RIP-DTR mice, with no observed mortality or overt signs of systemic toxicity. This contrasts sharply with STZ administration, which resulted in significant (p = 0.02) mortality in humanized mice (**Fig. 5D).** These findings highlight a key practical advantage of DT-based induction in this setting, particularly given the increased sensitivity of humanized models to systemic stressors.

Collectively, these findings demonstrate that DT provides a reliable and well-tolerated strategy for diabetes induction in huNBSGW RIP-DTR mice. The comparable incidence of hyperglycemia across the three female RIP-DTR cohorts, together with the absence of DT-associated toxicity, supports the utility of this model for studies integrating diabetes induction with human immune reconstitution.

### Humanized NBSGW RIP-DTR mice support evaluation of allogeneic immune responses to transplanted human islets and SCIs

To evaluate whether humanized NBSGW RIP-DTR mice can support immune-mediated responses to transplanted human grafts, graft function and graft-associated immune infiltration were assessed following renal subcapsular transplantation of human primary islets or SCIs. In a diabetic huNBSGW RIP-DTR mouse transplanted with human primary islets and analyzed 9 weeks after transplantation, fed BG levels progressively increased, whereas serum human C-peptide initially increased before declining at later time points, consistent with allograft rejection. (**Fig. 6A-F)**. Histological analysis identified residual graft tissue beneath the renal capsule (**Fig. 6A-B).** Immunofluorescent staining demonstrated residual insulin-positive cells within the graft, together with infiltration of human CD45^+^ leukocytes (**Fig. 6C**). Additional immune profiling identified graft-associated CD3^+^ T cells and CD19^+^ B cells (**Fig. 6D-E)**.

**Figure 6.**
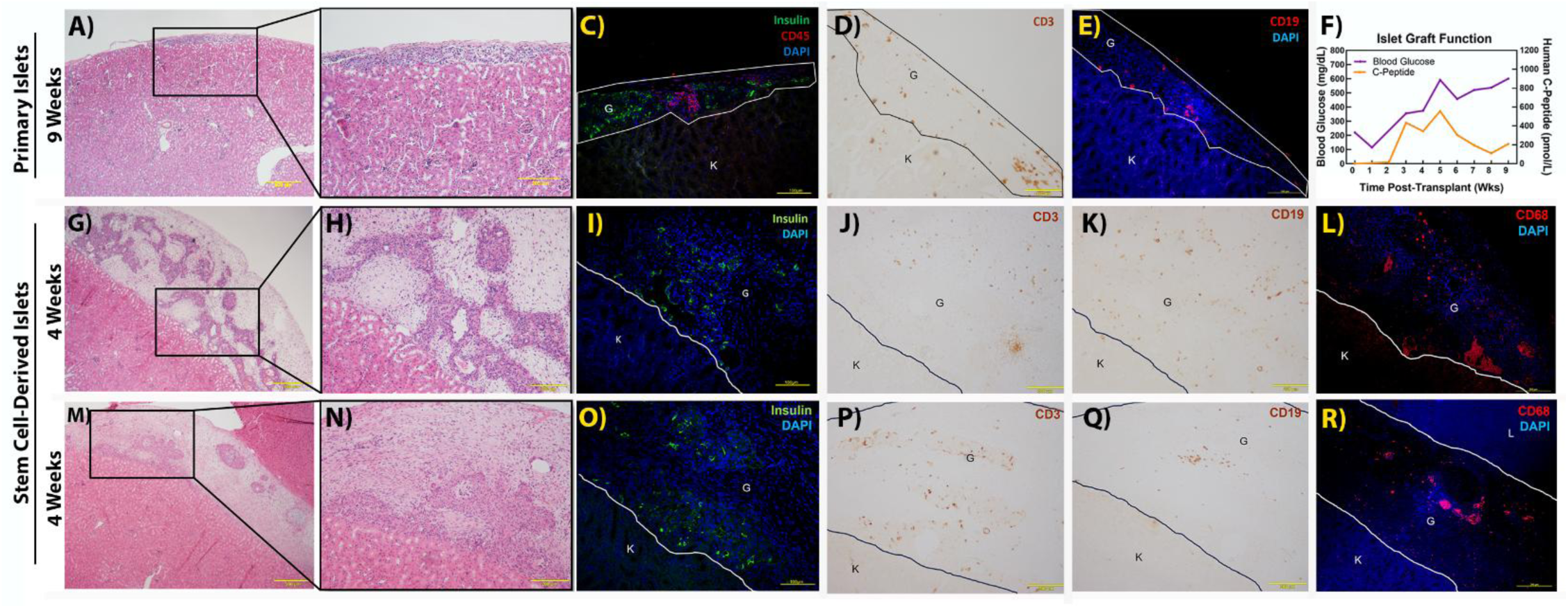
Immune cell infiltration following transplantation of human primary islets and stem cell-derived islets (SCIs) into humanized NBSGW RIP-DTR mice. Representative histologic analyses of renal subcapsular grafts following transplantation of **A-F)** human primary islets into a diabetic humanized (hu) NBSGW RIP-DTR mouse and **G-R)** stem cell-derived islets (SCIs) into huNBSGW mice and huNBSGW RIP-DTR mice. Primary human islet grafts were collected 9 weeks post-transplantation whereas SCI grafts were collected 4 weeks post-transplantation. **Primary human islet transplantation A-F)**. **A-B)** H&E staining showing residual graft tissue beneath the renal capsule, with the right panel showing higher magnification of the boxed region in **A**. **C)** Immunofluorescence (IF) staining for human insulin (green) and CD45^+^ leukocytes (red). **D)** Immunohistochemistry (IHC) for human CD3^+^ T cells. **E)** IF staining for human CD19^+^ B cells (red). **F)** Longitudinal fed BG and serum human C-peptide from the same representative mouse, showing progressive hyperglycemia accompanied by declining human C-peptide after an initial post-transplant increase, consistent with graft rejection. **Stem cell-derived transplantation (G-R).** Representative huNBSGW **(G-L)** and huNBSGW RIP-DTR (**M-R)** mice were evaluated 4 weeks after transplantation. **G-H** and **M-N)** H&E staining showing residual graft tissue beneath the renal capsule, with the right panel showing higher magnification of the boxed region. **I, O)** Immunofluorescence (IF) staining for human insulin (green). IHC for human **J, P)** CD3^+^ T cells, and for **K, Q)** CD19^+^ B cells. **L, R)** IF staining for CD68^+^ macrophages/monocytes. Outlined regions identify the graft (G) and kidney (K).

To determine whether this model could also be used to evaluate immune responses to SCIs, grafts were examined 4 weeks after transplantation into huNBSGW and huNBSGW RIP-DTR mice for comparative analysis. Residual SCI graft tissue was detected beneath the renal capsule in both recipients (**Fig. 6G-H, M-N)**, and insulin-positive cells were present within the graft regions (**Fig. 6I, O)**. Human immune-cell infiltration was detected in SCI grafts, including CD3^+^ T cells, CD19^+^ B cells, and CD68^+^ macrophage/monocyte-lineage cells (**Fig. 6J-L, P-R)**. Together, these representative findings showed allograft-associated human immune cell infiltration accompanied by evidence of graft rejection, supporting the feasibility of using huNBSGW RIP-DTR mice to evaluate immune responses to transplanted human islets and SCIs. Larger cohorts will be required to define the frequency and cellular composition of rejection.

## DISCUSSION

The NBSGW RIP-DTR mouse model addresses a major barrier in preclinical diabetes research: the lack of a single platform that combines controlled, low-mortality diabetes induction, with irradiation-free human immune reconstitution. By introducing the RIP-DTR transgene onto the NBSGW background, this model integrates DT-mediated β-cell ablation with the capacity for human hematopoietic engraftment without myeloablative conditioning, while additionally providing an extended experimental window for studying graft rejection.

A key requirement for the success of this approach is preservation of reliable diabetes induction. The present study demonstrates that DT-mediated β-cell ablation remains effective following transfer of the RIP-DTR transgene to the NBSGW background. Importantly, the optimized DT regimens identified in NSG RIP-DTR mice translated directly to the NBSGW RIP-DTR background without loss of induction efficiency or durability. These finding indicate that incorporation of the Kit*^W^*^41^ mutation does not interfere with the RIP-DTR-mediated β-cell targeting and supports the use of this model in studies requiring consistent and sustained hyperglycemia.

The β-cell specificity of the RIP-DTR system represents another important advantage of this model. Although STZ remains widely used for experimental diabetes induction, its utility is limited by strain-dependent variability, incomplete β-cell destruction, and off-target toxicity that can complicate interpretation of both metabolic and immune outcomes. ^3–5^ In contrast, DT selectively ablates β-cells while preserving neighboring endocrine cell populations, including α-cells, resulting in a more physiologically intact pancreatic islet. This selective targeting is particularly valuable for transplantation studies, where preservation of the recipient islet microenvironment allows immune-mediated graft injury to be evaluated without the confounding effects of widespread pancreatic damage. Moreover, reliable hyperglycemia was achieved using only nanogram quantities of DT, minimizing systemic exposure and reducing the risk of off-target toxicity. The improved tolerability of DT observed in this study, particularly in humanized mice, further supports its use in long term transplantation studies, where systemic stress could otherwise influence graft function and immune responses.

Beyond diabetes induction, the NBSGW RIP-DTR mouse supports functional human islet engraftment without compromising metabolic outcomes. Transplanted mice demonstrated sustained glycemic improvement, detectable human C-peptide, and glucose-responsive insulin secretion, indicating that the incorporation of the NBSGW background does not impair human islet engraftment and metabolic rescue. These findings establish that incorporation of the humanization-permissive background is compatible with β-cell replacement strategies and does not introduce a functional penalty at the level of graft performance.

Sex-associated trends in the onset of DT-induced hyperglycemia were observed in NSG RIP-DTR mice. Male mice showed a modestly earlier rise in BG following DT administration than females. These findings are consistent with previous reports demonstrating that sex can influence metabolic phenotypes in murine models.^25,26^ However, the present study was not designed or powered to define the mechanisms underlying these sex-associated differences. Importantly, sex did not alter identification of the optimized DT regimens or the central conclusion that NBSGW RIP-DTR mice retain efficient DT-mediated diabetes induction.

An important finding of this study is that the introduction of the RIP-DTR transgene did not impair irradiation-free human immune reconstitution, a notable feature of the NBSGW parental strain.^15, 16^. Human CD45^+^, CD3^+^, and CD19^+^ cell engraftment remained comparable between NBSGW RIP-DTR and parental NBSGW mice throughout the study, indicating that expression of the RIP-DTR allele did not adversely affect human hematopoietic engraftment or lymphoid development. This preservation of human immune reconstitution allowed diabetes induction and subsequent evaluation of human immune responses to occur within the same animals, providing a practical platform for longitudinal transplantation studies. Importantly, the extended survival and low mortality observed following DT administration enabled assessment of graft rejection over several weeks, a feature that is particularly valuable for studies of SCI maturation and immune-mediated graft loss.

The diabetes induction results obtained in humanized NBSGW RIP-DTR mice further support the feasibility of this integrated approach. Administration of only nanogram quantities of DT (15 ng total) induced hyperglycemia in humanized mice with an incidence comparable to that observed in non-humanized NBSGW RIP-DTR and NSG RIP-DTR mice, indicating that human immune reconstitution did not impair the efficiency of diabetes induction. Importantly, DT was well tolerated, with no observed mortality or overt signs of systemic toxicity, whereas STZ administration resulted in significantly increased mortality. These findings highlight that reliable diabetes induction can be achieved using very low doses of DT, providing an important practical advantage for long-term studies in humanized mice where systemic toxicity could otherwise confound immune and metabolic outcomes.

The humanized transplantation studies provide proof-of-feasibility that huNBSGW RIP-DTR can be used to evaluate human immune responses to transplanted human grafts. Representative primary human islet and SCI recipients demonstrated graft-associated multi-lineage immune-cell infiltration together with evidence of graft rejection, including progressive hyperglycemia, declining human C-peptide, and infiltration of human immune cells. Although these findings are based on representative cases, they demonstrate that the model supports longitudinal assessment of both graft function and immune-cell infiltration following transplantation. This capability provides a valuable platform for investigating mechanisms of allograft rejection and immune-cell trafficking and for preclinical evaluation of hypoimmune-gene edited SCIs and other strategies designed to improve graft survival. ^27–29^

Several limitations should be considered. First humanized mouse experiments are inherently variable due to donor-to-donor differences in human tissues and the stochastic nature of immune engraftment. Second, sex-associated differences in the onset of DT-induced hyperglycemia were observed and should be considered in future applications. Third, although DT is intended to selectively ablate murine β-cells in RIP-DTR mice, indirect effects of DT exposure on the broader graft microenvironment cannot be fully excluded. Finally, diabetes induction in humanized NBSGW RIP-DTR was slightly less penetrant than non-humanized mice, indicating that additional optimization may improve the efficiency of future study designs. These limitations outline areas for possible future refinement but do not diminish the utility of the model as an integrated experimental platform.

In summary, the NBSGW RIP-DTR model provides an integrated platform that combines controlled β-cell ablation, reversal of diabetes by human islet grafts, and compatibility with irradiation-free human immune reconstitution. This design enables more direct investigation of cadaveric and PSC-derived β-cell replacement strategies in the context of human immunity and reduces reliance on separate model systems. As such, the model represents a practical and translationally relevant tool for advancing preclinical studies of islet transplantation and immune-mediated graft responses.

## ACKNOWLEDGEMENTS

The authors would like to offer sincere thanks to the families who donated tissues, without which this study would not be possible. The graphical abstract was created using Illustrae.co. Human pancreatic islets were provided by the NIDDK-funded Integrated Islet Distribution Program (IIDP; RRID: SCR_014387) at City of Hope, NIH Grant # U24DK098085.

## Author Contributions

CSC: supervision, investigation, formal analysis, data curation, writing-original draft, writing-review and editing; DK: investigation, methodology, data curation, visualization, writing-original draft; EAS: investigation, methodology, data curation; LH: investigation, data curation, formal analysis; AMH: investigation, data curation, formal analysis; CL: visualization, investigation, writing-original draft; APK: investigation, data curation; KMG: investigation, data curation; CCS: investigation; AM: investigation; DMT: investigation; PC: investigation; KT: writing-original draft; EE: writing-original draft; BM: conceptualization; MEB: conceptualization, funding acquisition, methodology, supervision, formal analysis, writing-review and editing; SDS: conceptualization, funding acquisition, methodology, supervision, writing-review and editing; JSO: methodology, supervision, writing-review and editing.

## FUNDING

This work was, in part, supported by grants from the Breakthrough TID (BT1D) organization (3-SRA-2023-1419-S-B, 1-SRA-2016-168-S-B, and 1-PNF-2016-250-S-B to J.S.O.; and 2-SRA-2025-1662-S-B to M.E.B.), as well as by National Institutes of Health (NIH) grants R21-AI126419-01 (J.S.O), Department of Defense PR190621 (J.S.O.) F31 DK125021-01 (D.M.T.), NIH NIAID 75N93021C00004 (M.E.B.), the Wisconsin Alumni Research Foundation (WARF, M.E.B.). Additional support for this research was provided by the University of Wisconsin–Madison Office of the Vice Chancellor for Research and Graduate Education (OVCRGE) with funding from the Wisconsin Alumni Research Foundation (WARF) or OVCRGE/WARF. Lastly, data presented here were in part obtained through support from an NIH/NCATS UL1TR002373 award through the University of Wisconsin Institute for Clinical and Translational Research (UW ICTR to S.D.S and M.E.B). The content is solely the responsibility of the authors and does not necessarily represent the official views of Breakthrough TID, the National Institutes of Health, the Wisconsin Alumni Research Foundation, or the University of Wisconsin Institute for Clinical and Translational Research.

## CONFLICT OF INTEREST

J.S.O. is co-founder, chief scientific advisor, and holds stock equity of Regenerative Medical Solutions, Inc. and a consultant for Century Therapeutics ad GC Therapeutics. M.E.B is a founder of Allostasis Bio and a consultant for Taconic Biosciences. BEM is an employee and equity owner of Labcorp.

## Notes

### Summary of Updates

author affiliations updated and conflict of interest was updated

## References

1. Hogenes M, Huibers M, Kroone C, de Weger R. Humanized mouse models in transplantation research. Transplant Rev (Orlando). 2014;28(3):103–10. Epub 20140214. doi: 10.1016/j.trre.2014.02.002. PubMed PMID: 24636846.

2. Kenney LL, Shultz LD, Greiner DL, Brehm MA. Humanized Mouse Models for Transplant Immunology. Am J Transplant. 2016;16(2):389–97. Epub 20151120. doi: 10.1111/ajt.13520. PubMed PMID: 26588186; PMCID: PMC5283075.

3. Furman BL. Streptozotocin-Induced Diabetic Models in Mice and Rats. Curr Protoc. 2021;1(4):e78. doi: 10.1002/cpz1.78. PubMed PMID: 33905609.

4. Graham ML, Janecek JL, Kittredge JA, Hering BJ, Schuurman HJ. The streptozotocin-induced diabetic nude mouse model: differences between animals from different sources. Comp Med. 2011;61(4):356–60. PubMed PMID: 22330251; PMCID: PMC3155402.

5. Hosokawa M, Dolci W, Thorens B. Differential sensitivity of GLUT1- and GLUT2-expressing beta cells to streptozotocin. Biochem Biophys Res Commun. 2001;289(5):1114–7. doi: 10.1006/bbrc.2001.6145. PubMed PMID: 11741307.

6. Sudarshan MA, L.; Sharan, S. . Modeling Diabetes Mellitus Using Streptozotocin: Review Approach For Future Diabetic Research. IOSR-JBB. 2025;11(3):12–21. doi: 0.9790/ 264X-1103011321.

7. Ito M, Hiramatsu H, Kobayashi K, Suzue K, Kawahata M, Hioki K, Ueyama Y, Koyanagi Y, Sugamura K, Tsuji K, Heike T, Nakahata T. NOD/SCID/gamma(c)(null) mouse: an excellent recipient mouse model for engraftment of human cells. Blood. 2002;100(9):3175–82. doi: 10.1182/blood-2001-12-0207. PubMed PMID: 12384415.

8. King M, Pearson T, Rossini AA, Shultz LD, Greiner DL. Humanized mice for the study of type 1 diabetes and beta cell function. Ann N Y Acad Sci. 2008;1150:46–53. doi: 10.1196/annals.1447.009. PubMed PMID: 19120266; PMCID: PMC2620029.

9. Shultz LD, Lyons BL, Burzenski LM, Gott B, Chen X, Chaleff S, Kotb M, Gillies SD, King M, Mangada J, Greiner DL, Handgretinger R. Human lymphoid and myeloid cell development in NOD/LtSz-scid IL2R gamma null mice engrafted with mobilized human hemopoietic stem cells. J Immunol. 2005;174(10):6477–89. doi: 10.4049/jimmunol.174.10.6477. PubMed PMID: 15879151.

10. Herrera PL, Huarte J, Zufferey R, Nichols A, Mermillod B, Philippe J, Muniesa P, Sanvito F, Orci L, Vassalli JD. Ablation of islet endocrine cells by targeted expression of hormone-promoter-driven toxigenes. Proc Natl Acad Sci U S A. 1994;91(26):12999–3003. doi: 10.1073/pnas.91.26.12999. PubMed PMID: 7809163; PMCID: PMC45568.

11. Lee FT, Dangi A, Shah S, Burnette M, Yang YG, Kirk AD, Hering BJ, Miller SD, Luo X. Rejection of xenogeneic porcine islets in humanized mice is characterized by graft-infiltrating Th17 cells and activated B cells. Am J Transplant. 2020;20(6):1538–50. Epub 20200121. doi: 10.1111/ajt.15763. PubMed PMID: 31883299; PMCID: PMC7286695.

12. Thorel F, Nepote V, Avril I, Kohno K, Desgraz R, Chera S, Herrera PL. Conversion of adult pancreatic alpha-cells to beta-cells after extreme beta-cell loss. Nature. 2010;464(7292):1149–54. Epub 20100404. doi: 10.1038/nature08894. PubMed PMID: 20364121; PMCID: PMC2877635.

13. Bhagchandani P, Chang CA, Zhao W, Ghila L, Herrera PL, Chera S, Kim SK. Islet cell replacement and transplantation immunology in a mouse strain with inducible diabetes. Sci Rep. 2022;12(1):9033. Epub 20220531. doi: 10.1038/s41598-022-13087-3. PubMed PMID: 35641781; PMCID: PMC9156753.

14. Rongvaux A, Willinger T, Takizawa H, Rathinam C, Auerbach W, Murphy AJ, Valenzuela DM, Yancopoulos GD, Eynon EE, Stevens S, Manz MG, Flavell RA. Human thrombopoietin knockin mice efficiently support human hematopoiesis in vivo. Proc Natl Acad Sci U S A. 2011;108(6):2378–83. Epub 20110124. doi: 10.1073/pnas.1019524108. PubMed PMID: 21262827; PMCID: PMC3038726.

15. McIntosh BE, Brown ME, Duffin BM, Maufort JP, Vereide DT, Slukvin, II, Thomson JA. Nonirradiated NOD,B6.SCID Il2rgamma-/-Kit(W41/W41) (NBSGW) mice support multilineage engraftment of human hematopoietic cells. Stem Cell Reports. 2015;4(2):171–80. Epub 20150115. doi: 10.1016/j.stemcr.2014.12.005. PubMed PMID: 25601207; PMCID: PMC4325197.

16. McIntosh BE, Brown ME. No irradiation required: The future of humanized immune system modeling in murine hosts. Chimerism. 2015;6(1-2):40–5. Epub 20160512. doi: 10.1080/19381956.2016.1162360. PubMed PMID: 27171577; PMCID: PMC5063082.

17. Lan P, Tonomura N, Shimizu A, Wang S, Yang YG. Reconstitution of a functional human immune system in immunodeficient mice through combined human fetal thymus/liver and CD34+ cell transplantation. Blood. 2006;108(2):487–92. Epub 20060112. doi: 10.1182/blood-2005-11-4388. PubMed PMID: 16410443.

18. Brown ME, Zhou Y, McIntosh BE, Norman IG, Lou HE, Biermann M, Sullivan JA, Kamp TJ, Thomson JA, Anagnostopoulos PV, Burlingham WJ. A Humanized Mouse Model Generated Using Surplus Neonatal Tissue. Stem Cell Reports. 2018;10(4):1175–83. Epub 20180322. doi: 10.1016/j.stemcr.2018.02.011. PubMed PMID: 29576539; PMCID: PMC5998340.

19. Kalscheuer H, Danzl N, Onoe T, Faust T, Winchester R, Goland R, Greenberg E, Spitzer TR, Savage DG, Tahara H, Choi G, Yang YG, Sykes M. A model for personalized in vivo analysis of human immune responsiveness. Sci Transl Med. 2012;4(125):125ra30. doi: 10.1126/scitranslmed.3003481. PubMed PMID: 22422991; PMCID: PMC3697150.

20. Del Rio NM, Huang L, Murphy L, Babu JS, Daffada CM, Haynes WJ, Keck JG, Brehm MA, Shultz LD, Brown ME. Generation of the NeoThy mouse model for human immune system studies. Lab Anim (NY). 2023;52(7):149–68. Epub 20230629. doi: 10.1038/s41684-023-01196-z. PubMed PMID: 37386161; PMCID: PMC10935607.

21. Hogrebe NJ, Maxwell KG, Augsornworawat P, Millman JR. Generation of insulin-producing pancreatic beta cells from multiple human stem cell lines. Nat Protoc. 2021;16(9):4109–43. Epub 20210804. doi: 10.1038/s41596-021-00560-y. PubMed PMID: 34349281; PMCID: PMC8529911.

22. Ma SN, Yuan YH, Guo XR, Li DS. Subcapsular Implantation of Pancreatic Islets in Syngeneic, Allogeneic, and Xenogeneic Mice. Transplant Proc. 2016;48(8):2821–5. doi: 10.1016/j.transproceed.2016.06.045. PubMed PMID: 27788824.

23. Lemos JRN, Baidal DA, Poggioli R, Fuenmayor V, Chavez C, Alvarez A, Linetsky E, Mauvais-Jarvis F, Ricordi C, Alejandro R. Prolonged Islet Allograft Function is Associated With Female Sex in Patients After Islet Transplantation. J Clin Endocrinol Metab. 2022;107(3):e973–e9. doi: 10.1210/clinem/dgab787. PubMed PMID: 34727179; PMCID: PMC8852206.

24. Gannon M, Kulkarni RN, Tse HM, Mauvais-Jarvis F. Sex differences underlying pancreatic islet biology and its dysfunction. Mol Metab. 2018;15:82–91. Epub 20180530. doi: 10.1016/j.molmet.2018.05.017. PubMed PMID: 29891438; PMCID: PMC6066785.

25. Kim B, Park ES, Lee JS, Suh JG. Sex-specific differences in glucose metabolism and pancreatic function in streptozotocin-induced diabetic mice: The protective role of estrogen. Biochem Biophys Res Commun. 2025;775:152176. Epub 20250607. doi: 10.1016/j.bbrc.2025.152176. PubMed PMID: 40499498.

26. Mauvais-Jarvis F. Gender differences in glucose homeostasis and diabetes. Physiol Behav. 2018;187:20–3. Epub 20170824. doi: 10.1016/j.physbeh.2017.08.016. PubMed PMID: 28843891; PMCID: PMC5826763.

27. Sackett SD, Kaplan SJ, Mitchell SA, Brown ME, Burrack AL, Grey S, Huangfu D, Odorico J. Genetic Engineering of Immune Evasive Stem Cell-Derived Islets. Transpl Int. 2022;35:10817. Epub 20221205. doi: 10.3389/ti.2022.10817. PubMed PMID: 36545154; PMCID: PMC9762357.

28. Saha S, Haynes WJ, Seo J, Del Rio NM, Young EE, Zhang J, Holm AM, Pimentel M, Flannagan L, Huang L, Blashka W, Murphy L, Scholz MJ, Henrichs A, Suresh Babu J, Steill J, Kratz J, Stewart R, Kamp TJ, Brown ME. Diminished immune cell adhesion in hypoimmune ICAM-1 knockout human pluripotent stem cells. Nat Commun. 2025;16(1):7415. Epub 20250812. doi: 10.1038/s41467-025-62568-2. PubMed PMID: 40796566; PMCID: PMC12343971.

29. Reichman TW, Markmann JF, Odorico J, Witkowski P, Fung JJ, Wijkstrom M, Kandeel F, de Koning EJP, Peters AL, Mathieu C, Kean LS, Bruinsma BG, Wang C, Mascia M, Sanna B, Marigowda G, Pagliuca F, Melton D, Ricordi C, Rickels MR, Group V-FS. Stem Cell-Derived, Fully Differentiated Islets for Type 1 Diabetes. N Engl J Med. 2025;393(9):858–68. Epub 20250620. doi: 10.1056/NEJMoa2506549. PubMed PMID: 40544428.

