## Supplementary material for "The NBSGW RIP-DTR Mouse: An Integrated Platform for Diabetes Induction, Human Immune Reconstitution and Transplantation Studies": Online supplemental materials

**Supplementary Methods**

**Animals**

NOD.Cg-*Prkdc^scid^ Il2rg^tm1Wjl^* Tg(Ins2-HBEGF)6832Ugfm/SzJ (NSG RIP-DTR; strain no. 027976) and NOD.Cg-*Kit^W-41J^ Tyr^+^ Prkdc^scid^ Il2rg^tm1Wjl^*/ThomJ (NBSGW; strain no. 026622) mice were obtained from the Jackson Laboratory.

**Generation of NBSGW RIP-DTR Mice**

To generate NBSGW RIP-DTR mice, NSG RIP-DTR mice were crossed with NBSGW mice to produce F1 progeny. Animals carrying both the *Ins2-HBEGF/RIP-DTR* transgene and the *Kit^W41^* allele were identified and selected for continued breeding. Selected animals were backcrossed to the NBSGW strain for 10 generations to establish a homozygous *Kit^W41^* background while retaining the RIP-DTR transgene.

Genotyping was performed using PCR-based genotyping (Transnetyx, Cordova, TN) to detect the *Ins2-HBEGF* (RIP-DTR transgene), *Kit^W41^* mutation, *Prkdc^scid^* allele and *Il2rg-null* allele. Genotyping was conducted at each generation during backcrossing to confirm allele status. Experimental mice were required to be homozygous for the *Kit^W4/W41^* mutation, positive for the RIP-DTR transgene, and homozygous for both *Prkdc^scid^* and *Il2rg-null* alleles before inclusion. Coat color was monitored across generations as an additional indicator of strain convergence.

**Generation of Humanized NeoThy Mice**

NeoThy humanized mice were generated as previously described^1,2^. Briefly, human thymic fragments (approximately 1 mm^3^) were implanted beneath the kidney capsule. Immediately thereafter, mice received 0.5-1.5 x 10^5^ allogeneic CD34^+^ human umbilical cord hematopoietic stem/progenitor cells (HSPCs) via tail-vein injection. CD34^+^ HSPCs were isolated by magnetic bead selection (Miltenyi Biotec, Bergisch Gladback, Germany). Peripheral blood was collected at 8, 12, and 15-16 weeks after humanization for flow cytometric analysis.

**Diphtheria Toxin (DT) Preparation**

Lyophilized diphtheria toxin (List Biological Laboratories, #150) was reconstituted at 1mg/ml in sterile distilled water, aliquoted, and stored at –20°C. Working dilutions were prepared fresh in sterile 0.9% NaCl immediately before use. DT was administered intraperitoneally, and blood glucose (BG) was monitored longitudinally to assess onset, penetrance, and durability of diabetes. Multiple DT dosing regimens, including single doses of 10 ng, 15 ng (15 ng x 1), or >30 ng, and 5 ng administered daily for 3 consecutive days (5 ng x 3), were first evaluated in NSG RIP-DTR mice to assess induction kinetics, penetrance, and toxicity. To mitigate severe hyperglycemia during DT long-term durability studies, LinBit insulin pellets (LinShin Canada, Inc., Ontario, Canada) were implanted 2 wk post-DT when BG exceeded 300 mg/dL.

**Stem Cell-Derived Islet Differentiation**

Pluripotent HUES8 (Harvard University, Cambridge, MA, USA) human embryonic stem cells were maintained under feeder-free conditions and differentiated into SCIs using a six-stage differentiation protocol described by Hogrebe et al.^3^. Differentiation cultures were maintained in suspension throughout endocrine specification and maturation with media changes performed according to the published protocol. SCI clusters were harvested upon completion of Stage 6 and used for renal capsule transplantation. Differentiation quality was monitored throughout the protocol by morphology and routine assessment of pancreatic lineage marker expression. All reagents and concentrations were used as reported by Hogrebe et al.

**Human Islet Transplantation**

NBSGW RIP-DTR and humanized NBSGW RIP-DTR (huNBSGW RIP-DTR) mice underwent renal subcapsular transplantation using aseptic conditions. Under isoflurane anesthesia, a flank incision was made to expose the right kidney, and a small incision was created in the renal capsule. Human pancreatic islets (2,000 IEQ) or HUES8 stem cell-derived islets (SCIs; 10,000 IEQ) were loaded into polyethylene (PE)-50 tubing attached to a Hamilton syringe and transplanted beneath the kidney capsule. The kidney was returned to the peritoneal cavity, and the incision was closed with absorbable sutures and tissue adhesive. Age- and sex-matched NSG RIP-DTR mice were transplanted in parallel as comparator controls. Fed BG was measured twice weekly. Peripheral blood was collected before transplantation and every 2 weeks thereafter for human C-peptide analysis. During episodes of sustained hyperglycemia, mice received exogenous insulin therapy consisting of SQ long-acting insulin glargine (1-2 U; Sanofi-Aventis) and/or LinBit sustained release implants (LinShin Canada, Toronto, Canado). LinBit implants release approximately 0.1 U insulin per day for >30 days. At study completion, mice underwent intraperitoneal glucose tolerance testing (GTT), followed by kidney graft harvest for histology.

**Metabolic Assessment and C-Peptide Measurement**

Peripheral blood (100-150 µl) was collected by retro-orbital venipuncture into serum separator tubes containing aprotinin. Samples were allowed to clot for 30 min at room temperature and centrifuged at 11,000 rpm for 5 minutes at 4°C. Serum was collected and stored at -80°C until analysis.

Human serum C-peptide was measured using an Ultrasensitive C-peptide ELISA kit according to the manufacturer’s instructions (Mercodia, Uppsala, Sweden). The assay is specific for human C-peptide with negligible cross-reactivity to mouse C-peptide (<0.1%). Samples were run in duplicate and absorbance was measured at 450 nm using a FlexStation 3 plate reader (Molecular Devices, San Jose, CA).

**Intraperitoneal Glucose Tolerance Testing**

Mice were fasted overnight (≥12 hours) with free access to water. A sterile 30% D-glucose solution (Sigma G-7528) was administered intraperitoneally (i.p.) at 2 g/kg body weight. BG was measured at 0-, 15-, 30-, 60-, 90-, and 120-min. Blood samples were collected at baseline and 60 min for C-peptide analysis.

**Histology and Immunofluorescence**

At the study endpoint, transplanted kidneys were harvested, fixed in 4% paraformaldehyde for 24h, paraffin-embedded, and sectioned at 5 µm. For immunofluorescence, sections were deparaffinized, rehydrated, and subjected to heat-induced antigen retrieval in 10 mmol/L sodium citrate buffer (pH 6.0). Sections were permeabilized with 0.05% Triton X-100, blocked in 10% bovine serum albumin, and incubated with primary antibodies, followed by species-appropriate fluorophore-conjugated secondary antibodies for 40 min at room temperature in the dark. When multiple primary antibodies were used, species-appropriate combinations and sequential staining were used to minimize cross-reactivity. Nuclei were counterstained with DAPI. Negative control sections were processed identically except that the primary antibodies were omitted. Detailed antibody information is provided in **Supplementary Table 2**.

**Imaging**

Fluorescence images were acquired using a Zeiss Axiovert 200M microscope equipped with an AxioCam HR camera and controlled by AxioVision software (Release 4.8; both from (Carl Zeiss Microscopy GmbH, Oberkochen, Germany). Brightfield images were acquired using a Nikon Eclipse E600 microscope (Nikon Corporation, Tokyo, Japan) equipped with an Olympus DP73 camera controlled by CellSens imaging software (Olympus Life Science, Tokyo, Japan).

**Flow Cytometric Analysis of Human Engraftment**

Peripheral blood was collected by retro-orbital sampling into heparinized tubes. Red blood cells were removed using ACK lysis buffer. Cells were washed in FACS Buffer and stained with fluorophore-conjugated antibodies against human and mouse CD45 (pan-leukocytes), and human CD3 (T-cells), and CD19 (B-cells). Dead cells were excluded before analysis. Flow cytometry was performed using a BD Aria III, BD Accuri cytometers (BD Biosciences, San Jose, CA), or CytoFLEX (Beckman Coulter, Indianapolis, IN) and analyzed using FlowJo v10 (BD Biosciences, Ashland, OR). Human immune engraftment was expressed as the percentage of human CD45^+^ leukocytes among total CD45^+^ leukocytes (defined as human CD45^+^ plus mouse CD45^+^ cells)^4-5^. B cell and T-cell frequencies were quantified as percentages of human CD45^+^ cells unless otherwise indicated. Compensation was performed using single stained compensation beads (BD), and gates were established using unstained controls. Detailed antibody information is provided in **Supplementary Table 2**.

**Supplementary Table 1. Human islet donor characteristics**

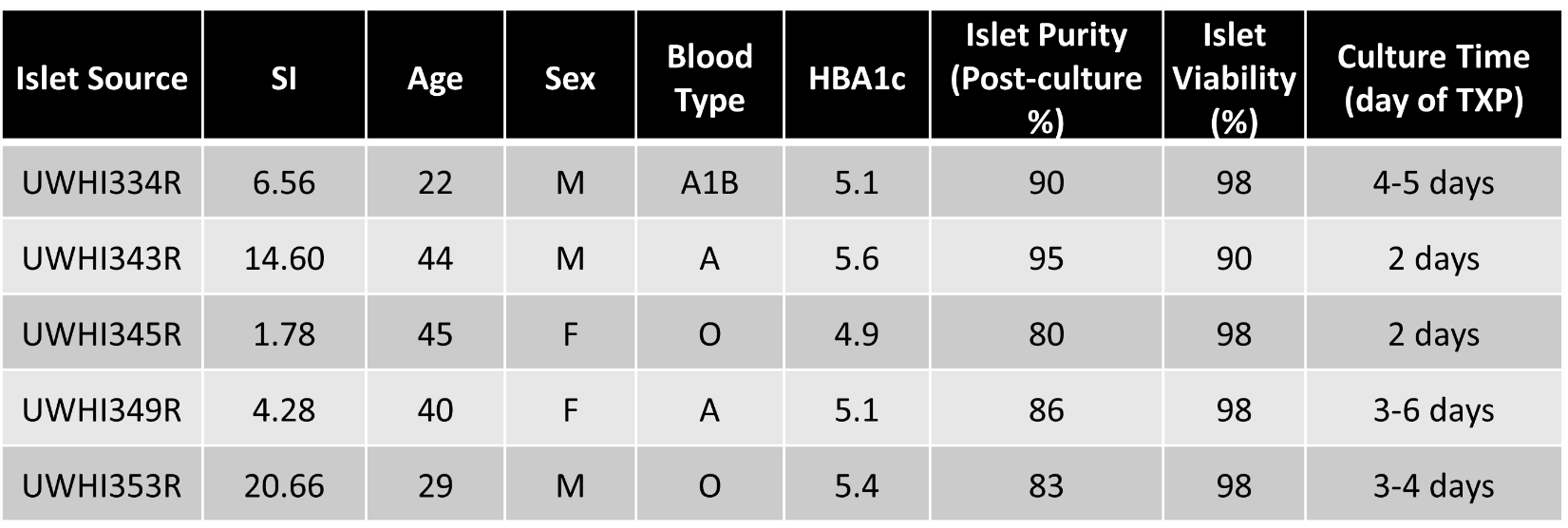

**Supplementary Table 2. Antibody information used for immunofluorescence.**

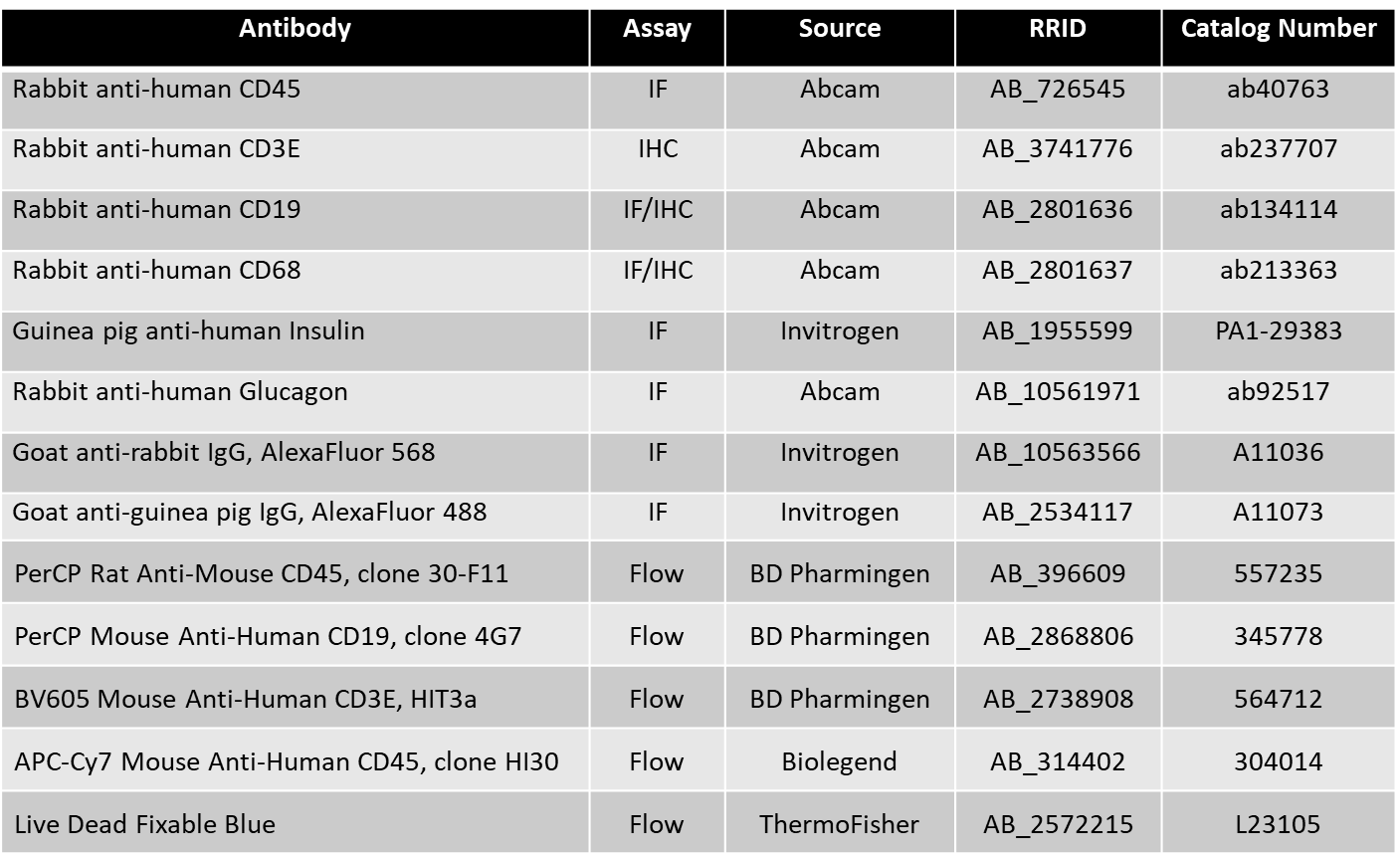

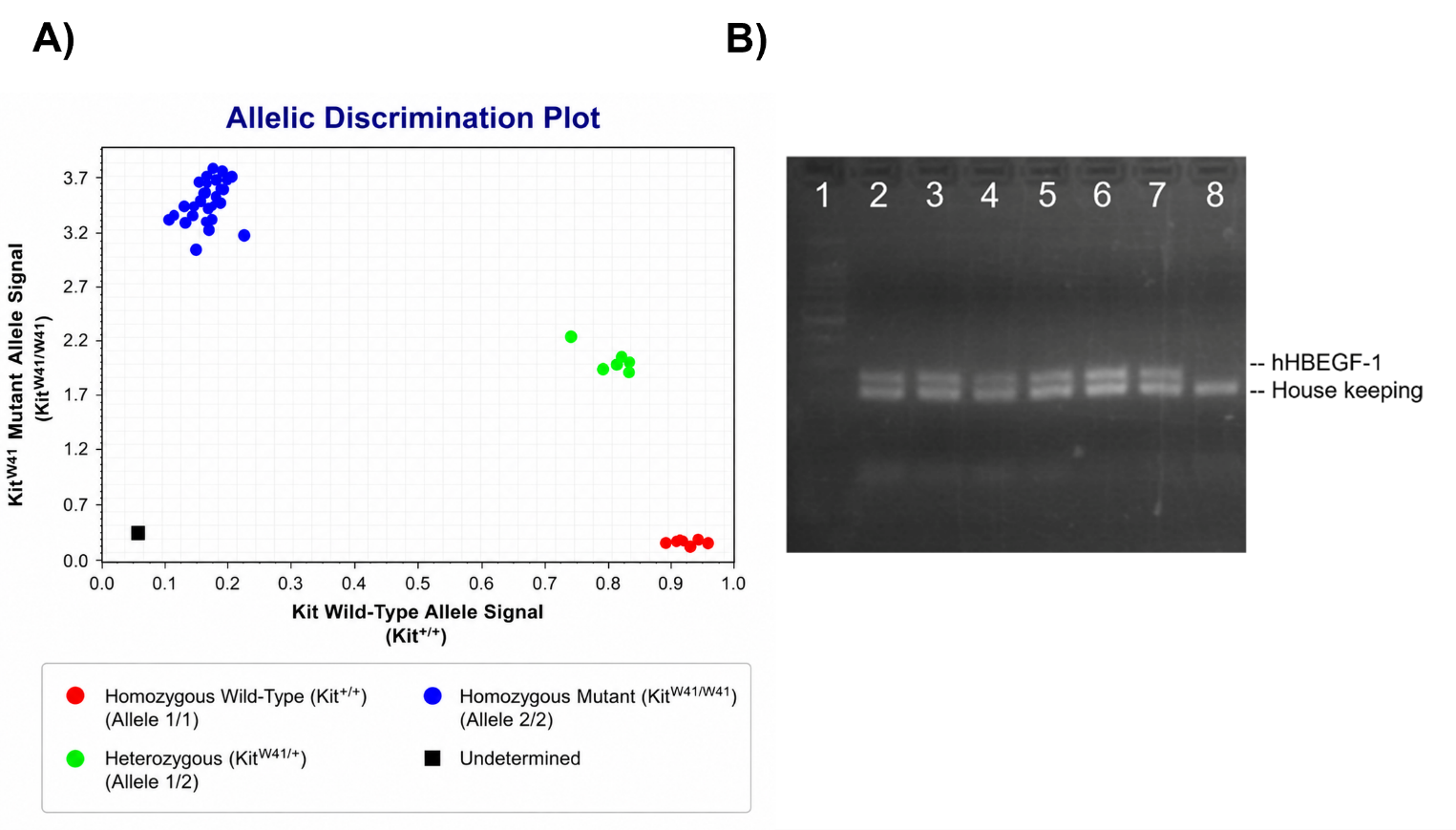

**Supplementary Figure 1: Genotype verification of NBSGW RIP-DTR mice by allelic discrimination and transgene analysis.** Representative PCR-based genotyping results confirming the genetic integrity of the NBSGW RIP-DTR strain. **A)** Allelic discrimination plot from a TaqMan PCR-based assay targeting the *Kit^W41^* locus, showing clear separation of homozygous *Kit^W41/W41^* (blue), heterozygous *Kit^W41/+^* (green), and wild type^(+/+)^ (red) genotypes. Each data point represents an individual mouse. Experimental NBSGW RIP-DTR mice cluster exclusively within the homozygous *Kit^W41/W41^* population, confirming fixation of the NBSGW-defining allele. **B)** Representative PCR gel images demonstrating presence of the human *HBEGF-1* (Ins2-DTR) transgene in experimental mice. Lane 1 contains the molecular weight ladder. Lanes 2-7 correspond to individual mice positive for the hHBEGF-1 (Ins2-DTR) transgene, while lane 8 represents a mouse lacking the transgene and serves as the negative control. Only mice meeting all inclusion criteria were used for subsequent experiments.

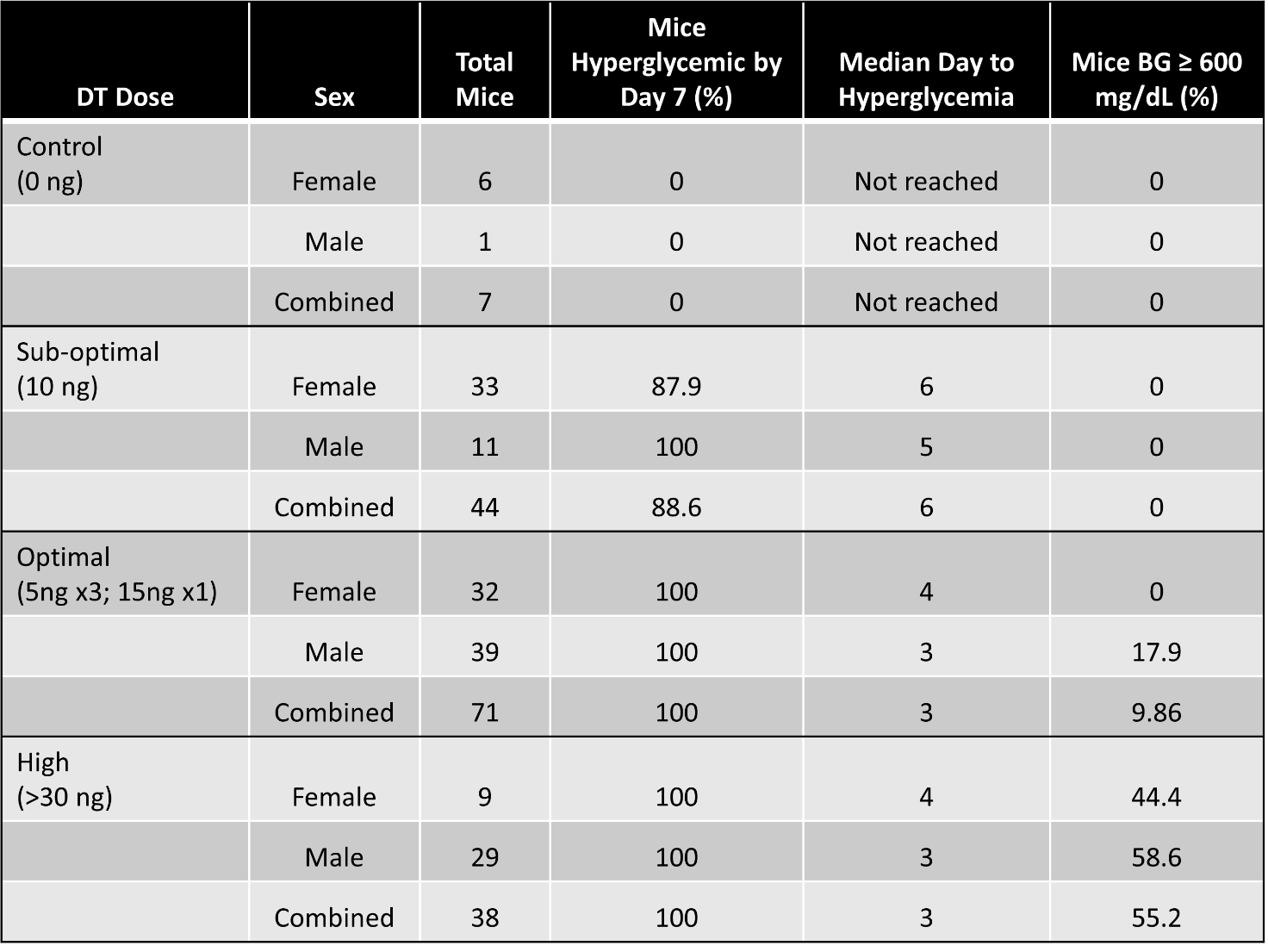

**Supplementary Table 3. Summary of diphtheria toxin (DT) dose optimization for hyperglycemia induction in NSG RIP-DTR mice. A)** Mice were grouped according to DT dosing regimen and satisfied by sex or analyzed as a combined cohort. Hyperglycemia was defined as BG ≥ 300 mg/dL, and the percentage of hyperglycemic mice was determined by day 7 following DT administration. The median day to hyperglycemia and the percentage of mice reaching the upper detection limit of the glucometer (BG ≥ 600 mg/dL) are reported as measures of hyperglycemia onset and severity, respectively. Data are presented as percent hyperglycemia and/or time where indicated.

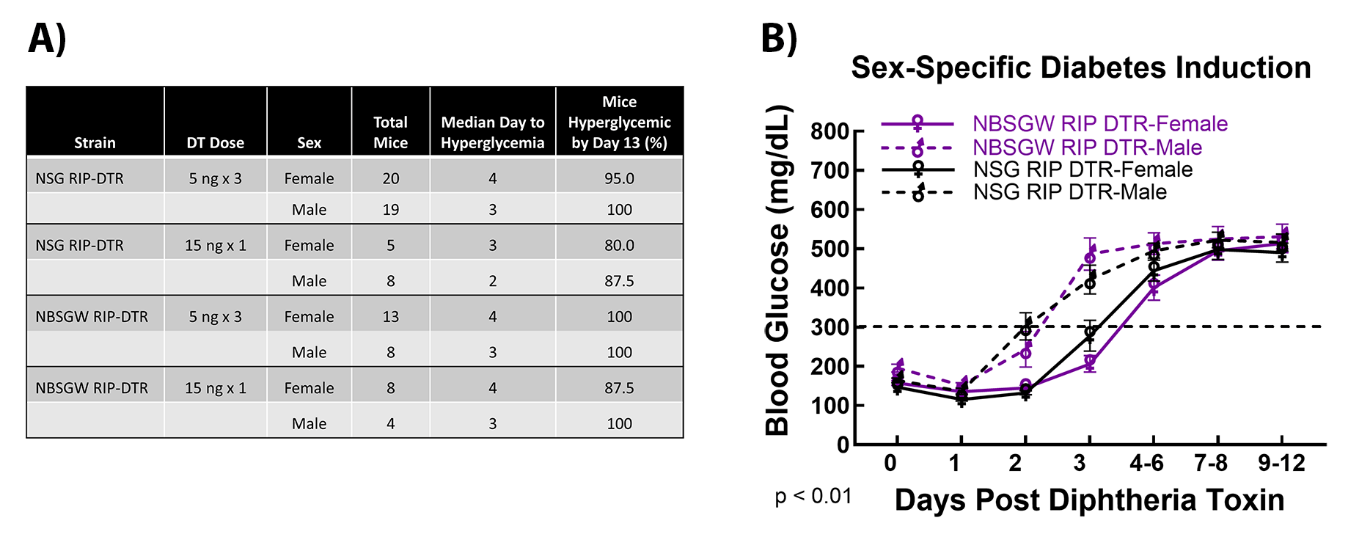

**Supplementary Figure 2. Sex-specific diabetes induction following diphtheria toxin (DT) administration.** **A)** Percent hyperglycemia by day 13 stratified by strain, DT regimen, and sex. **B)** Longitudinal BG trajectories stratified by strain and sex. Mice receiving the 5 ng x 3 and 15 ng x 1 DT regimens were combined for analysis. BG trajectories differed significantly among groups over time (mixed-effects model, p < 0.01) Post-hoc analysis demonstrated an earlier rise in BG in males than females, with differences most evident between days 2 and 3. Data are presented as mean ± SEM. Statistical analysis was performed using mixed effects models (REML) and Tukey’s post hoc correction for multiple comparisons.

1.	Del Rio, N.M., Huang, L., Murphy, L., Babu, J.S., Daffada, C.M., Haynes, W.J., Keck, J.G., Brehm, M.A., Shultz, L.D., and Brown, M.E. (2023). Generation of the NeoThy mouse model for human immune system studies. Lab Anim (NY) *52*, 149-168. 10.1038/s41684-023-01196-z.

2.	Brown, M.E., Zhou, Y., McIntosh, B.E., Norman, I.G., Lou, H.E., Biermann, M., Sullivan, J.A., Kamp, T.J., Thomson, J.A., Anagnostopoulos, P.V., and Burlingham, W.J. (2018). A Humanized Mouse Model Generated Using Surplus Neonatal Tissue. Stem Cell Reports *10*, 1175-1183. 10.1016/j.stemcr.2018.02.011.

3.	Hogrebe, N.J., Maxwell, K.G., Augsornworawat, P., and Millman, J.R. (2021). Generation of insulin-producing pancreatic beta cells from multiple human stem cell lines. Nat Protoc *16*, 4109-4143. 10.1038/s41596-021-00560-y.

4.	Saito, Y., Ellegast, J.M., Rafiei, A., Song, Y., Kull, D., Heikenwalder, M., Rongvaux, A., Halene, S., Flavell, R.A., and Manz, M.G. (2016). Peripheral blood CD34(+) cells efficiently engraft human cytokine knock-in mice. Blood *128*, 1829-1833. 10.1182/blood-2015-10-676452.

5.	Saito, Y., Ellegast, J.M., Rafiei, A., Song, Y., Kull, D., Heikenwalder, M., Rongvaux, A., Halene, S., Flavell, R.A., and Manz, M.G. (2016). Peripheral blood CD34+ cells efficiently engraft human cytokine knock-in mice. Blood *128*, 1829-1833. 10.1182/blood-2015-10-676452.
